# Elucidating the combined toxicity of aflatoxin B1 and fumonisin B1 on HepG2 cells based on respirometry and transcriptome analyses

**DOI:** 10.1101/2023.01.19.524737

**Authors:** Xiangrong Chen, Mohamed F. Abdallah, Charlotte Grootaert, Filip Van Nieuwerburgh, Andreja Rajkovic

## Abstract

Aflatoxin B1 (AFB1) and fumonisin B1 (FB1) are two toxic mycotoxins widely found in food contaminants, and known for their hepatotoxicity in human. However, their combined toxicity still needs to be deeply investigated especially for their harmful effect. Therefore, the current work aimed at investigating the (combined) effect of AFB1 and FB1 on mitochondrial and glycolytic activity of HepG2 cell line, a well-recognized *in vitro* model system to study liver cell function. In our previous work, we studied the impact of a short term exposure to different doses of AFB1, FB1, and their binary mixture (MIX) on the bioenergetic status of HepG2 cells. Seahorse respirometry analysis revealed that the co-exposure, especially at high doses (8 µg/mL for AFB1 and 160 µg/mL for FB1), is more toxic as a result of more inhibition of all parameters of mitochondrial respiration. RNA transcriptome sequencing showed that the p53 signaling pathway, which is a major orchestrator of mitochondrial apoptosis, was differentially expressed. Moreover, the co-exposure has significantly downregulated Cx I, Cx II, Cx III, and Cx IV genes, which represent the onset of the suppressed mitochondrial respiration in HepG2 cells. It was found that FB1 is contributed more to the MIX effects than AFB1.⍰

**Environmental Implication:** Aflatoxin B1 (AFB1) and fumonisin B1 (FB1) are two main mycotoxins that frequently (co-)contaminate maize and maize-based ingredients in several parts of the world. Both toxins are well-known for their hepatotoxicity in humans as the liver is their main target organ. However, the combined toxicity of AFB1 and FB1 still needs to be deeply investigated especially for their effect on cellular respiration. In this study, we proved that a binary mixture of AFB1 and FB1 is more toxic on mitochondrial respiration, and disrupted the p53 signaling pathway to induce apoptosis, which promised a novel insight of hazardous materials-induced hepatic damage.

## 1. Introduction

Recent advances in mycotoxin analysis showed that the co-occurrence of several mycotoxins in a single food or crop is more common than the occurrence of a single mycotoxin (Palumbo et al. 2020). However, most of the available knowledge regarding the toxicity of mycotoxins in human is limited to the study of a single mycotoxin, and little is known about the interaction of a mycotoxin mixture in the biological systems. Such interaction could be additive, antagonistic, or synergistic, which may alter the toxic outcomes (Alam et al. 2022). Currently, the number of mycotoxins in food is expected to be much more than 400, varying in their chemical structures and hence their toxicities (Decleer et al. 2018; Palumbo et al. 2020; Wu et al. 2014). In the present work, we have selected two mycotoxins (aflatoxin B1 and fumonisin B1) which have been found to co-occur in different food samples, especially maize, collected from Africa, America, Asia, and Europe (Chen et al. 2021; Du et al. 2017; Palumbo et al. 2020). Besides, aflatoxin B1 (AFB1) and fumonisin B1 (FB1) are considered among the most toxic fungal secondary metabolites. The International Agency for Research on Cancer (IARC) classified AFB1 as a group 1 carcinogen due to the sufficient evidence of causing liver cancer in humans, while FB1 is classified as a class 2B carcinogen as the evidence of causing cancer is limited (IARC 2012). Other toxic effects of AFB1 and FB1 also include hepatotoxicity, nephrotoxicity, and embryotoxicity, immunotoxicity (Chen et al. 2022).

Mitochondria are critical cellular organelles that make adenosine triphosphate (ATP) appropriately in response to the cellular energy demands, hence known as the powerhouse of the cell (Vyas et al. 2016). Besides, these organelles perform many roles, including the generation of reactive oxygen species (ROS) and regulation of cell signaling and cell death. Due to their highly abundance in hepatocytes, they have been recognized as a key mediator in hepatotoxicity. Such toxicity could be induced by the loss of mitochondrial function creating a mitochondrial metabolic gridlock, such as the inhibition of mitochondrial respiration (Prakash et al. 2022). The *in vitro* hepatotoxicity of AFB1 and FB1 has been reported in many studies using the HepG2 cells as the preferred liver model (Abdul and Chuturgoon 2021; Chen et al. 2022; Singto et al. 2020). Oxidative stress, inflammation, and mitochondrial dysfunction by targeting ROS, DNA, p53, and other signaling pathways have been documented as toxic mechanism of AFB1 (Li et al. 2022). Similarly, FB1 induced hepatotoxicity by inhibiting sphingolipids biosynthesis and triggering massive production of ROS have been also reported (Sheik Abdul and Marnewick 2020).

Based on these studies, it was found that both AFB1 and FB1 could damage mitochondrial function to cause hepatotoxicity. Currently, there are few reports on the use of *in vitro* systems for the analysis of AFB1 and FB1 mixtures, especially the effect on mitochondria. Therefore, the aim of the study reported here was to evaluate the interaction of AFB1 and FB1 in HepG2 cells. We investigated the impact of AFB1 and FB1 as well as their combination (binary mixture) on the mitochondrial and glycolytic activities using seahorse extra-cellular flux analysis that decipher bioenergetic phenotype, and by transcriptome analysis using RNA (Illumina) sequencing to better understand functional biology underlying observed phenotypic mitochondrial function signatures.

## 2. Materials and Methods

### 2.1. Chemical reagents

The mycotoxin FB1 (Cas. No. 116355-83-0; 99 % purity) was obtained from Sigma (USA), while AFB1 (Cas. ALX-630-093-M005; >98 % purity) was purchased from Enzo Life Sciences (Belgium). Stock solutions (10 mg/mL) of FB1 and AFB1 were prepared in dimethyl sulfoxide (DMSO) at stored at −20 1, while working solutions were freshly prepared in a cell growth medium at different concentrations and combinations of FB1 and AFB1 (**Table 1**). Dulbecco’s modified Eagle’s medium (DMEM) supplemented with GlutaMAX™, a mixture of penicillin/streptomycin, and non-essential amino acids (NEAA) were all purchased from Thermo Life Technologies (Merelbeke, Belgium). Fetal Bovine Serum (FBS) was supplied from VWR (Leuven, Belgium). Trypsin-EDTA 0.05 % from Thermo Fisher Scientific (Merelbeke, Belgium). Phosphate buffer saline (PBS) with or without Ca^2+^ and Mg^2+^ were obtained from Westburg (Leusden, Netherlands).

**Table 1.** Different concentrations of AFB1 and FB1 as well as their combinations used in the current work.

| Concentrations | AFB1<br>(µg/mL) | FB1<br>(µg/mL) | MIX |  |
| --- | --- | --- | --- | --- |
|  |  |  | AFB1 (µg/mL) | FB1 (µg/mL) |
| Low | 0.5 | 10 | 0.5 | 10 |
| Middle | 2 | 40 | 2 | 40 |
| High | 8 | 160 | 8 | 160 |

### 2.2. Cell culture and mycotoxin exposure

HepG2 (human hepatocellular carcinoma) cells were obtained from the American Type Culture Collection (ATCC, Manassas, VA, USA) and cultured in DMEM medium (Gibco™, GlutaMAX™) containing 4.5 g/LD-glucose and pyruvate, and externally supplemented with FBS (10 %), mixture of penicillin/streptomycin (1 %) and NEAA (1 %). Cells were grown in T-75 (75 cm^2^) polystyrene cell culture flasks (Thermo Life Technologies, Merelbeke, Belgium) in a humidified chamber with 10 % CO_2_ at 37 ⍰ and 95 % air atmosphere at constant humidity. The growth culture medium was changed every 2-3 days and cell morphology was regularly checked by visual inspection with phase-contrast microscopy (Leica DMIC, Leica Microsystem GMbH, Wetzlar, Germany). When the degree of confluence reaches approximately 80 % (every 4-6 days), cells were subcultured to maintain the rapid growth of the cells. Consequently, HepG2 cells were gently rinsed with prewarmed PBS for a few seconds, detached from the flask with 4 mL of pre-warmed trypsin-EDTA 0.05 % for 2-3 min, and seeded in a new T-75 flask (ratio 1:5).

HepG2 cells were exposed to three different concentrations of AFB1 (0.5, 2, and 8 µg/mL) and FB1 (10, 40, and 160 µg/mL) to mimic three different scenarios of exposure (low, middle and high). Furthermore, we applied three combinations (low-low, middle-middle, and high-high) as a binary mixture (MIX) of AFB1 and FB1 (**Table 1**). These concentrations were applied based on the estimated exposure data derived from measuring the average of urinary biomarkers of AFB1 (0.5 µg/mL) and FB1(10 µg/mL) in humans, which were considered here in the current work as low exposure scenario (Chen et al. 2022; Meneely et al. 2018). This level was increased four folds to have a middle exposure scenario and eight folds as high exposure scenario to investigate the potential toxicity.

### 2.3. Cytotoxicity endpoint measurements (MTT, ROS, and MMP)

The tetrazolium salt (MTT) (Life Technologies Corporation, Eugene, OR, USA) assay, which is based on the cellular conversion of 3-(4,5-dimethylthiazol-2-yl)-2,5-diphenyltetrazolium bromide into formazan, was performed to determine the cell viability after the exposure to AFB1 and FB1 (Chen et al. 2022). Briefly, HepG2 cells were seeded before the treatment in 96-well plates at a density of 20,000 per well for 24 h at 37 ⍰ in a sterilized incubator with a humidified atmosphere of 10 % CO_2_ to allow cell adhesion. FB1 and AFB1 were diluted with DMEM at different concentrations and 200 µL were added. The applied toxic doses of AFB1, FB1, and their mixture (MIX) are shown in **Table 1**. MTT was measured using SpectraMax™ Microplate Reader (Molecular Devices, Berkshire, UK), as described in the literature (Chen et al., 2022). Levels of ROS and MMP (mitochondrial membrane potential) were measured to reflect the toxic effect of AFB1 and FB1 on HepG2 cells. ROS generation and MMP were measured using a fluorescent probe (2’, 7’-dichlorodihydrofluorescein diacetate (DCFH-DA)) (Sigma-Aldrich) and the fluorescent probe Tetramethylrhodamine ethyl ester (TMRE) (Sigma-Aldrich), respectively, as described in literature (Chen et al., 2022). Cells were seeded at a density of 50,000 per well in a black 96-well plate for 24 h, and incubated with FB1 and AFB1 at different concentrations for another 24 h (**Table 1**). ROS and MMP were measured using SpectraMax™ Microplate Reader (Molecular Devices, Berkshire, UK), and the exact excitation and emission wavelengths were followed as described in the literature (Chen et al., 2022).

### 2.4. HepG2 bioenergetic analysis using Seahorse Extracellular Flux Analyzer (total ATP production, glycolysis, and mitochondrial respiration)

The Seahorse XF96 Analyzer instrument (Agilent Seahorse Bioscience, CA, USA), and the related consumables (plates, cartridges, and inhibitor kits) were used to measure total ATP production, glycolysis, and mitochondrial respiration according to the manufacture instructions. In brief, the assay medium was prepared by supplementing Seahorse XF Base medium (pH 7.4) with a specific combination of 10 mM glucose (100X stock, Agilent), 1 mM pyruvate (100X stock, Agilent), and 2 mM L-glutamine (Sigma). At first, cells were harvested from the T-75 flasks and seeded into a Seahorse 96-well XF Cell Culture microplate (Agilent Seahorse Bioscience, CA, USA) in 80 µL of the culture medium at a density of 20,000/well. The optimal cell density was previously determined and is part of laboratory SOPs for different cell types. The cells were incubated at 37°C in a pre-sterilized incubator with atmosphere containing 10 % CO_2_ and 95 % constant humidity for 24 h. Next, HepG2 cells were treated with the same doses of AFB1, FB1, and MIX as shown in **Table 1**.

In parallel, a Seahorse XF Sensor Cartridge was hydrated one day before running the XF Assay by filling each well of the XF Utility Plate with 200 µL of sterile water. On the day of analysis, the sterile water was replaced by seahorse XF calibrant solution. The hydrated cartridge was for 24h maintained in an incubator at 37 °C without CO_2_ to remove CO_2_ from the media that may interfere with measurements by altering the pH. Cells washing and measurement cycles were performed following our established protocol (Chen et al., 2022). Preformulated and optimized Seahorse specific real-time ATP rate assay kit, glycolysis stress test kit, and Mito stress test kit (all from Agilent) were used to measure total ATP production, glycolysis, and mitochondrial respiration, respectively. The accurate concentrations and volumes for each inhibiting compound used in each kit are described in **Table 2**. Seahorse Wave Controller Software version 2.6.3 (Agilent Seahorse Bioscience, CA, USA) was used to operate and control the Seahorse XF96 Analyzer instrument. After the measurements were done, data were exported for processing and analysis (see **Data processing and analysis**). Normalization was performed by fixing the cells using sulforhodamine B dye (Sigma-Aldrich, Co., St. Louis, MO, USA) as described before (Chen et al., 2022).

**Table 2.**
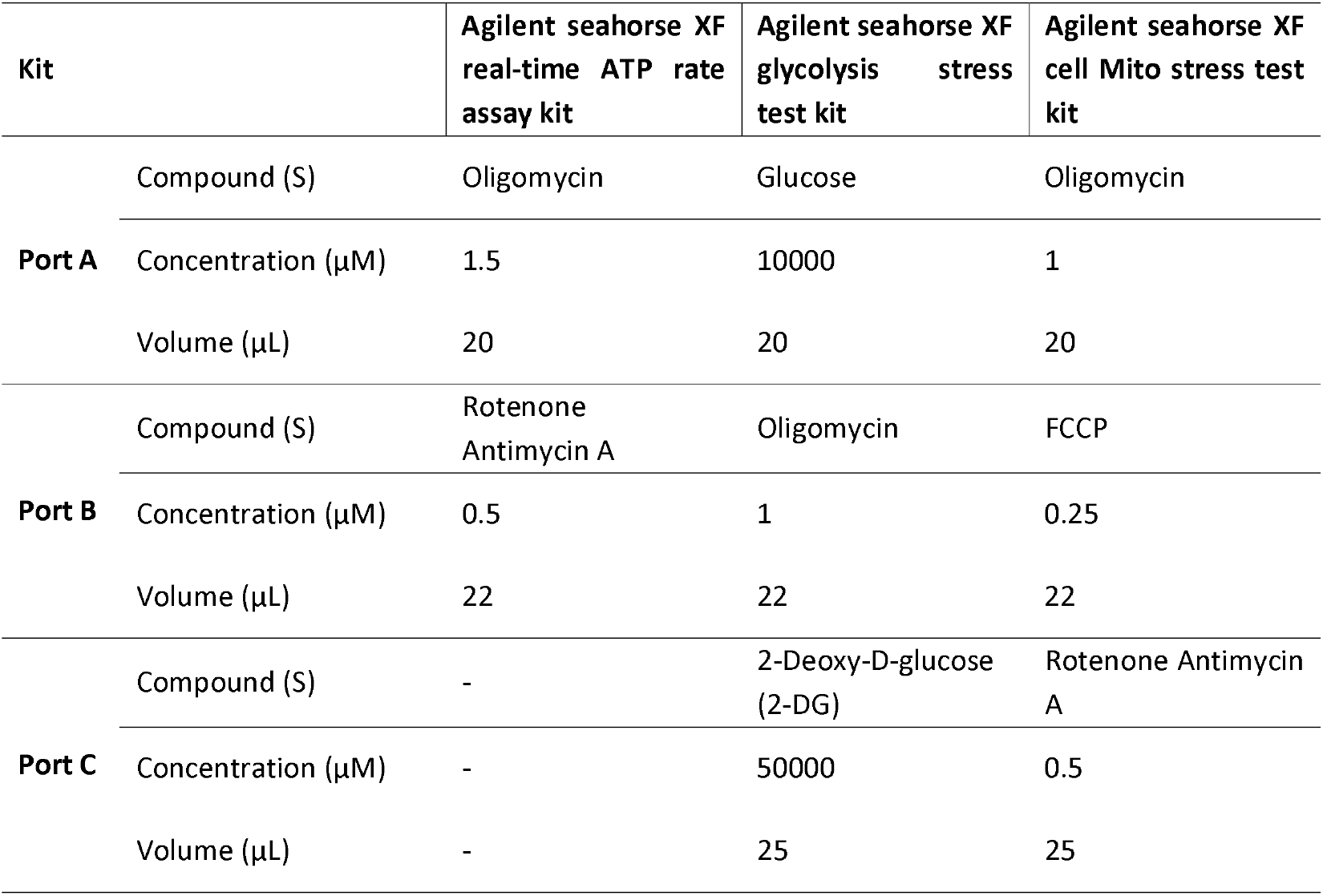
Required concentrations and volumes of each inhibitor per assay.

### 2.5. Transcriptome analysis (RNA isolation, processing and sequencing)

The complete set of RNA transcripts of HepG2 was studied at high concentrations of AFB1 (8 µg/mL) and FB1 (160 µg/mL) as well as their binary combination (high MIX) as these concentrations and combination induced toxic effect on mitochondria according the seahorse extra-cellular flux analysis. Transcriptome analysis including RNA isolation, processing and sequencing was conducted according to previously established protocols (Degroote et al. 2021). The experiment was repeated independently five times with identical conditions to provide at least five biological replicates for each time point and treatment. In summary, HepG2 cells were cultivated as previously mentioned but in six well-plates while keeping the same cell density. After 24 h of exposure to the mycotoxins under study, the growth media were completely removed. To harvest the cells, PBS (one mL) of was added to each well and the adherent cells were collected by cell scrapers (Greiner Bio-One, Vilvoorde, Belgium) and transferred into two mL Eppendorf tubes. After centrifugation for two min at 8000 g, the supernatant was discarded and the RNA extraction was performed with the RNeasy® Mini Kit (Qiagen, Hilden, Germany) following the instruction handbook. In short, 600 µL lysis Buffer RLT was applied to the pellets, 600 µL 70 % ethanol was added, and the total volume was transferred to an RNeasy Mini spin column placed in a collection tube to be centrifuged at 8000 g for 15 sec. The flow through is discarded, and the column is washed with Buffer RW1 and Buffer RPE, before eluting the RNA in 40 µL RNase-free water. A Bioanalyzer RNA 6000 Nano assay (Agilent Technologies, CA, USA) was used to measure RNA quality, providing a RIN (RNA Integrity Number) value. All the samples had an RNA integrity number (RIN) value above nine. RNA from each sample was quantified using the ‘Quant-it ribogreen RNA assay’ (Life Technologies, Grand Island, NE, USA) and 500 ng RNA was used to prepare an Illumina sequencing library using the QuantSeq 30 mRNA-Seq Library Prep Kit (Lexogen, Vienna, Austria) according to manufacturer’s protocol with 14 enrichment polymerase chain reaction (PCR) cycles. An average of 9.0 × 10^6^ ± 1.8 × 10^6^ and 11.6 × 10^6^ ± 1.0 × 10^6^ reads were generated.

### 2.6. Data processing and analysis

Excel© for Microsoft Office 365 (Microsoft Corporation, Redmond, USA) was used to normalize all data, except for transcriptome data. SPSS software package (SPSS Statistics 27, USA) was used for the statistical evaluation. Comparisons between the untreated control and different FB1 and AFB1 treatments within each mitochondrial parameter (basal respiration, maximal respiration, ATP production, proton leak, non-mitochondrial respiration, and spare respiratory capacity) were performed using a one-way analysis of variance (ANOVA) test followed by Tukey HSD multiple-comparison test as a post hoc analysis to identify the sources of detected significance (p < 0.05). Data are presented as mean ± standard deviation (SD). Analysis for differential gene expression was performed using the edgeR’s (40) quasi-likelihood method between 2 conditions, only including genes that were expressed at a counts-per-million (cpm) above 1 in at least 5 samples. Genes were considered significantly differential if they had a false discovery rate (FDR) < 0.05, as well as a fold change of at least 2. Gene set enrichment analysis (GSEA) was performed using the GAGE R package, based on the Kyoto encyclopedia of genes and genomes (KEGG) pathways provided by this package. Genes and KEGG pathways of interest were selected based on their impact on diabetes pathology, hepatic fat synthesis, and energy metabolism. Significance thresholds |log(FC)| >1 and FDR <0.05 were set in performing heatmaps; |log2FC| = 1 and P = 0.05 were set in performing volcano plots based on edgeR analysis. Finally, the effects of mycotoxins on pathways were also calculated and displayed in the heat map. The heat map expresses the magnitude enrichment analysis was performed using the GSEA software (v4.2.3) and Molecular Signatures Database (MSigDB) Hallmark Gene Signatures. Gene sets were considered significantly enriched when q-value FDR < 0.05, and normalized enrichment scores (NES) were used for further calculations of e.g. z-scores.

## 3. Results

### 3.1. Cytotoxicity of AFB1, FB1, and their combination after measuring MTT, ROS, and MMP

Treatment of HepG2 cells with either AFB1 or FB1 or their binary combination (MIX) resulted in a concentration-dependent increase in intracellular ROS, and induction of MMP disruption and therefore cell damage. As shown in **Figure 1**, a combination of AFB1 and FB1 at low concentration (low MIX) had no inhibitory effect on cellular viability in comparison to the individual effect imposed by either AFB1 or FB1 treatment. While a binary mixture of AFB1 and FB1 at middle and high concentrations (middle and high MIX) led to a significant loss of cell viability, which was between 9-24 % and 3-7 % higher compared to the loss of cell viability caused by AFB1 and FB1 alone, respectively. Significant increases in intracellular ROS levels were detected in case of exposure to AFB1 (high concentration) and FB1 (middle and high concentration) as well as the three levels of combinations (low, middle, and high MIX). The increase in ROS levels with a combination (MIX) was between 5-19 % more than the ROS levels with AFB1 and between 18-29 % higher than those levels detected with FB1. As depicted in **Figure 1**, the MMP disruption in HepG2 cells exposed to high concentrations of AFB1 (8 µg/mL) and FB1 (160 µg/mL) as well as their binary combination (high MIX) showed a slight but significant (p<0.05) dose-dependent MMP decrease. This decrease in MMP levels was about 12% for AFB1 and FB1 alone, and about 16% for the MIX, compared to the MMP from the untreated control cells. These results demonstrate that the applied doses of AFB1 and FB1 reduce cell viability and MMP and induce the generation of more intracellular ROS in HepG2 cells. By comparing the toxic impact of AFB1 or FB1 and their MIX in each exposure scenario (low, middle, and high), the binary combination (MIX) shows a trend to stronger effects in all cytotoxicity endpoints, although this was not always statistically different than the toxic effect caused by exposure to the individual AFB1 or FB1.

**Figure 1.**
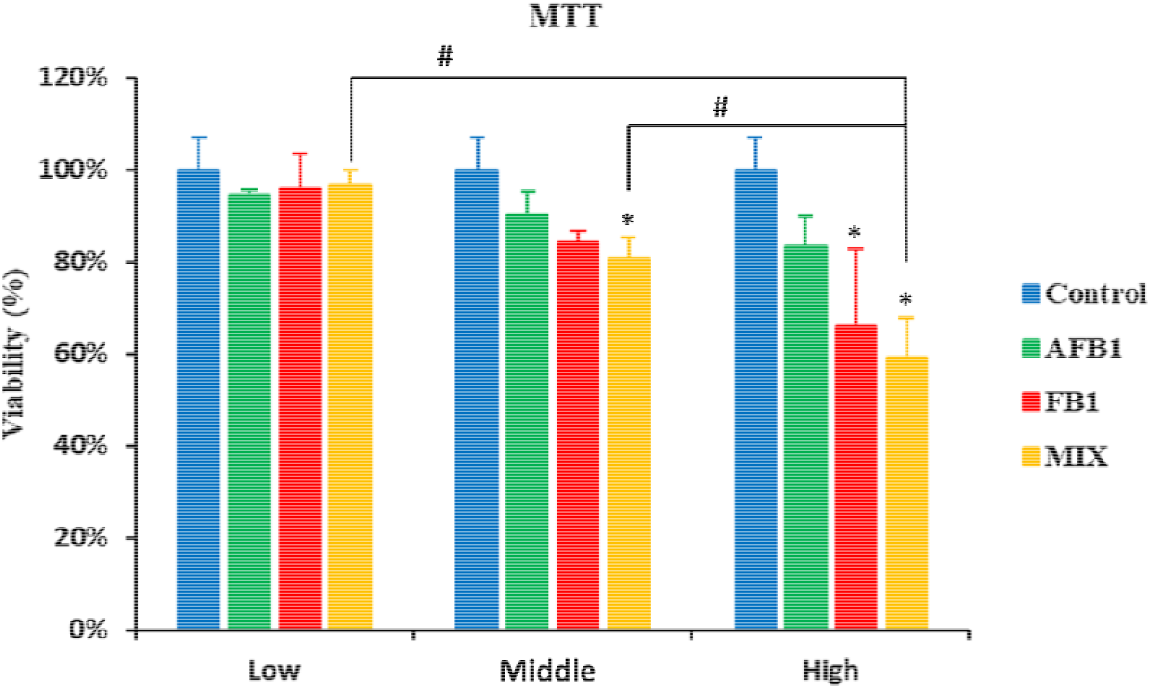
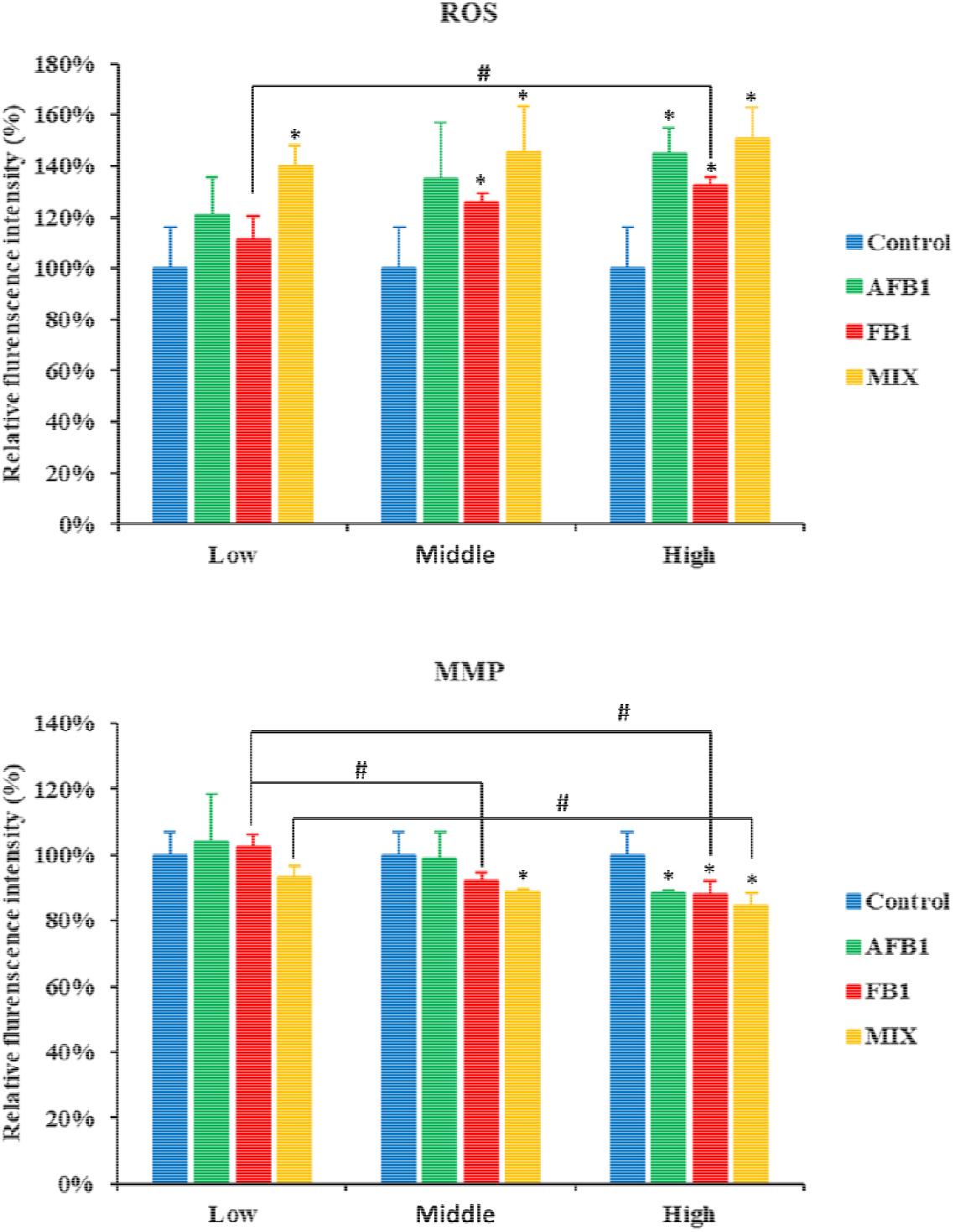
Effect of FB1, AFB1, and MIX on cell viability (MTT), intracellular reactive oxygen species (ROS), and mitochondrial membrane permeability (MMP) in HepG2 cells after 24 h exposure. Data (at least three well-replicates) are expressed as mean ± standard deviation. Significance compared to the mycotoxin-free condition is labeled by *: p < 0.05 according to a one-way ANOVA with Dunnett’s post hoc test. Significance compared to the same mycotoxin per concentration is labeled by #: p < 0.05 according to a one-way ANOVA with Tukey HSD multiple-comparison post hoc test.

### 3.2. Impact of AFB1, FB1 and their combination on HepG2 bioenergetics

#### 3.2.1 Total ATP production

To investigate the impact of AFB1 and FB1 as well as their combination on the total ATP production derived from glycolysis and mitochondrial respiration via oxidative phosphorylation (OXPHOS), the Seahorse XF Real-Time ATP Rate assay was used. As depicted in **Figure 2**, the total ATP production was inhibited in the three scenarios of exposure (low, middle, and high) either after individual treatment of AFB1 or FB1 or their binary combination (MIX) treatment in a concentration-dependent manner, which were statistically significant (p < 0.05). Interestingly, the exposure to AFB1 or FB1 or their combination (MIX) at the highest levels of exposure significantly shifted the balance or the contribution ratio of glycolysis versus OXPHOS for the total ATP yield to be more relying on the ATP generation via OXPHOS. After the exposure to MIX (low and middle level), the inhibition of the ATP production is situated between the effects of the single toxin treatments, thereby suggesting interactions at the energy-providing pathway level. In contrast, upon exposure to high MIX, it showed the strongest decrease (but not significant compared the individual exposure to toxins) in total ATP production, and caused a significant shift compared to the AFB1 and FB1 condition(s) from glycolytic to mitochondrial ATP production of 34 % that is the difference of OXPHOS ATP (67 %) and glycolysis (33 %). In various cancer cells such as HepG2 cells, glycolysis is enhanced and OXPHOS capacity is diminished. In the current work, the observed shifts in the contribution ratio between glycolysis and OXPHOS for the total ATP provide an unfavorable environment for cell growth (Zheng 2012).

**Figure 2.**
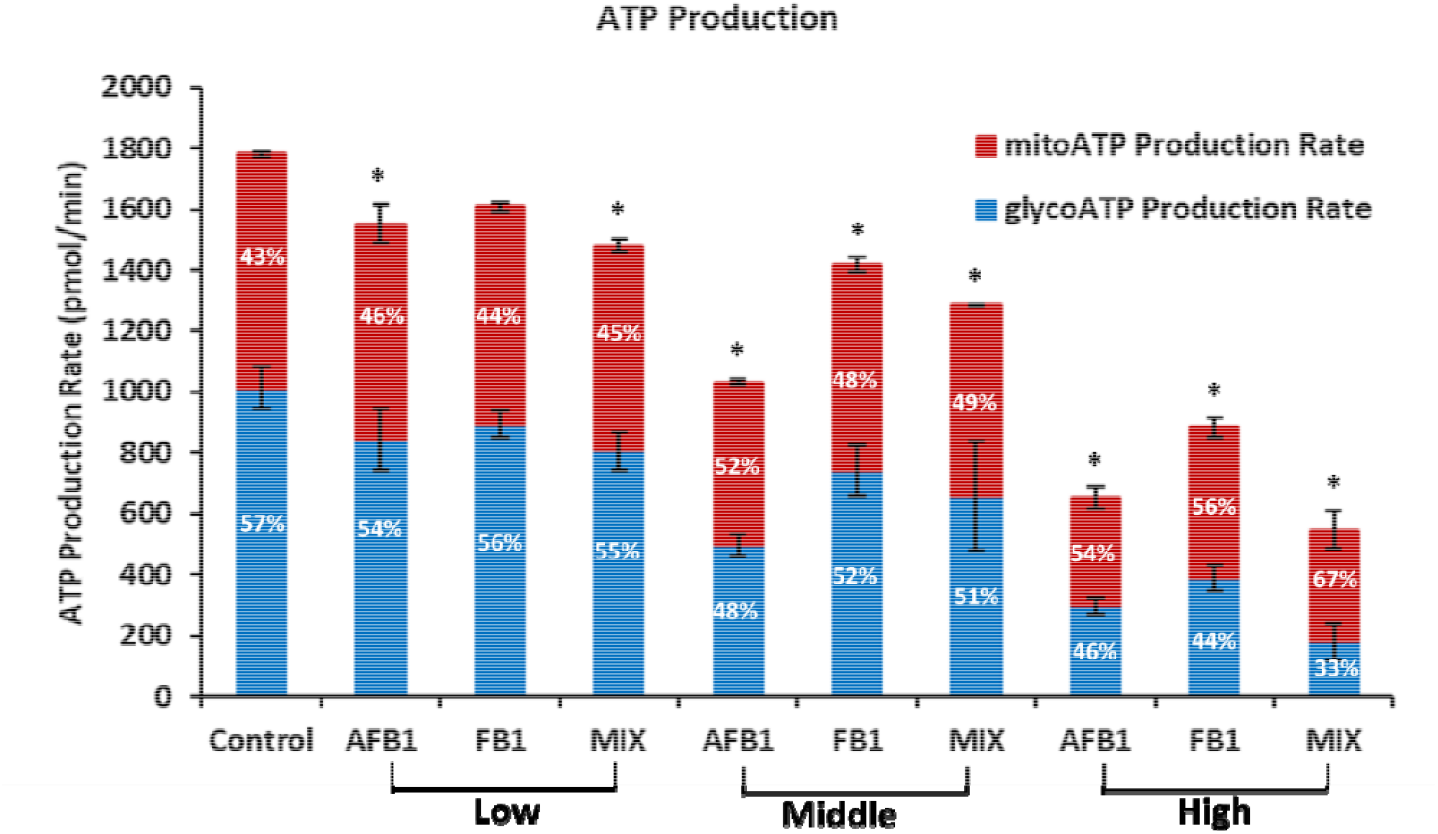
Effect of AFB1, FB1, and MIX on mitochondrial and glycolytic ATP production rates in HepG2 cells after 24 h exposure. Data (at least four technical replicates) are expressed as mean ± standard deviation. *: p < 0.05 indicates significantly different results compared to the untreated condition control by one-way ANOVA with Dunnett’s post hoc test.

#### 3.2.2. Glycolytic pathway for energy production in HepG2 cells

Basically, the glucose is converted into pyruvate (referred to as glycolysis) and then converted to lactate in the cytoplasm or CO_2_ and water in the mitochondria. The conversion of glucose to pyruvate, and subsequently lactate, results in net production and extrusion of protons into the extracellular medium. This extrusion of protons results in the acidification of the medium surrounding the cell. The extracellular acidification rate (ECAR) is related to the lactate secretion, which in turn is directly related to the glycolytic flux. Therefore, the glycolytic rate was measured as ECAR by the glycolytic rate using the Seahorse XF96 Analyzer. In the current work, the glycolytic activity of HepG2 cells was assessed following real-time changes in ECAR levels. After a 24 h treatment with either AFB1 or FB1 or their binary combination (MIX), there was a reduction in the glycolysis, glycolytic reserve, and glycolytic capacity (p < 0.05), except for the low exposure level (10 µg/mL) of FB1 treatment (**Figure 3**). When HepG2 cells were exposed to a combination of the two toxins (high MIX), the levels of glycolysis, glycolytic reserve, and glycolytic capacity were decreased in comparison to the individual treatment of AFB1 or FB1, however, these decreases were not statistically significant. These results demonstrate that high MIX (8 µg/mL for AFB1 and 160 µg/mL for FB1) might have a more disruptive effect on the glycolysis. However, the significant interaction of AFB1 and FB1 on the negative effect on glycolytic activity was not observed.

**Figure 3.**
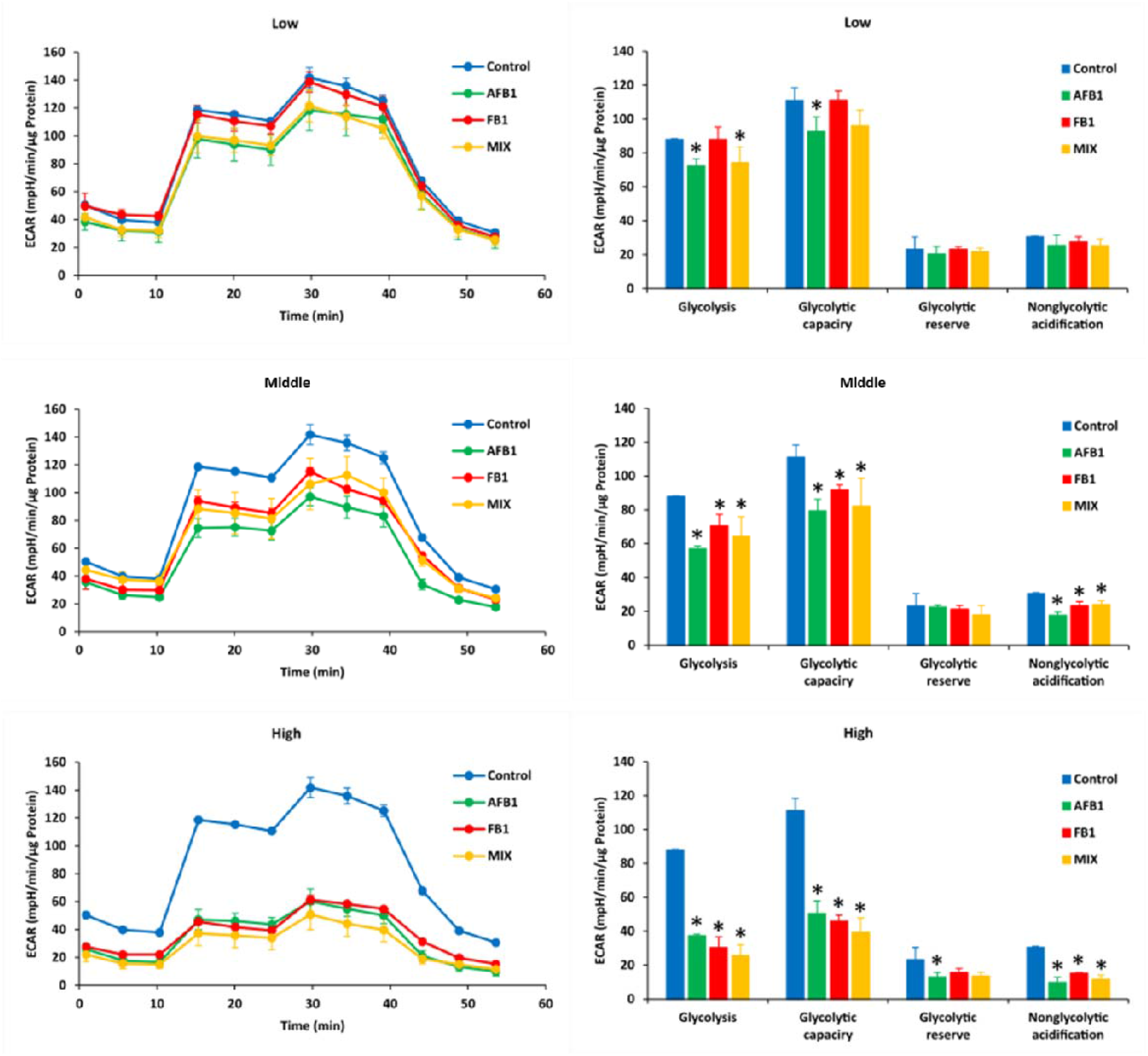
Effect of AFB1, FB1, and MIX on extracellular acidification rate (ECAR) (left) and different glycolytic parameters (right) in HepG2 cells after 24 h exposure. Data (at least four technical replicates) are expressed as mean ± standard deviation. Mean values with different symbols (*: compared to control) within each glycolytic parameter indicate significant differences (p < 0.05) among different treatments according to one way ANOVA test followed by the Tukey HSD multiple-comparison test as a post-doc analysis.

#### 3.2.3. Mitochondrial respiration pathway for energy production in HepG2 cells

In parallel, the effect of AFB1 and FB1 mixture on the ability of HepG2 cell to regulate mitochondrial respiration in comparison to the single exposure to AFB1 or FB1 was examined. In this regard, Oxygen Consumption Rate (OCR) is used as an indicator of mitochondrial respiration. An inhibition was observed in the mitochondrial activity of HepG2 cells after the exposure to three levels (low, middle, and high) of AFB1 or FB1 or their combination (MIX) (**Figure 4**). Especially the exposure to AFB1 or FB1 or their binary combination (MIX) at the high levels of exposure decreased the basal respiration, maximal respiration, ATP production, and proton leak (p < 0.05), and increased the spare respiratory capacity (p < 0.05) compared to untreated control. Moreover, when HepG2 cells were exposed to a combination of the two toxins (high MIX), all these mitochondrial parameters were significantly decreased in comparison to the individual treatment of high AFB1 (8 µg/mL) or high FB1 (160 µg/mL). However, these significant decreases were not always present in the other two exposure scenarios (low and middle) between comparisons of the individually toxic impact of AFB1 or FB1 with their binary combination (MIX). These results demonstrate that high MIX (8 µg/mL for AFB1 and 160 µg/mL for FB1) might cause more disruption of the mitochondrial metabolism, and a significant changes in mitochondrial dysfunction seem to be attributed to the toxic outcome of the interaction between AFB1 and FB1.

**Figure 4.**
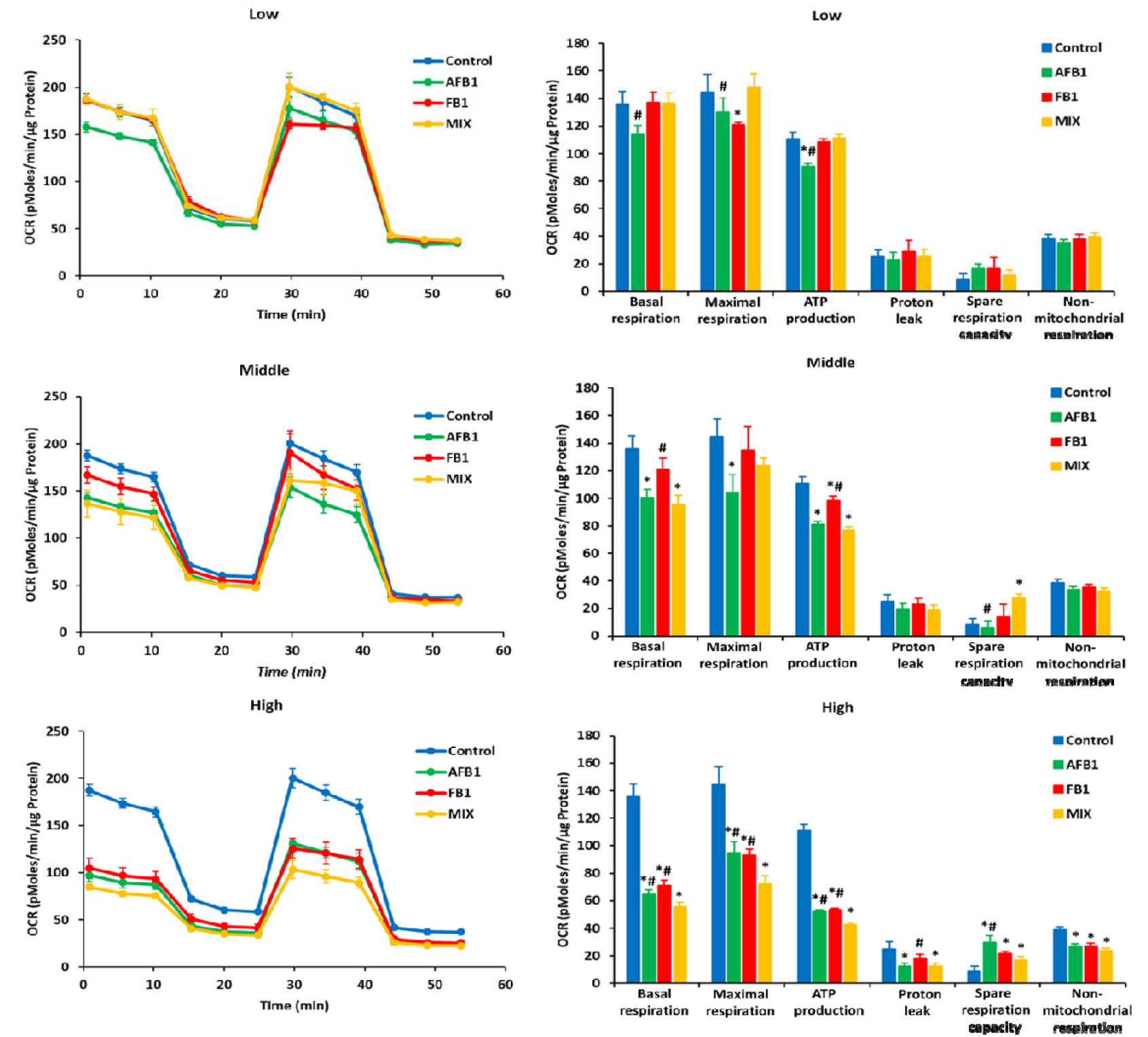
Effect of AFB1, FB1, and MIX on oxygen consumption rate (OCR) (left) and different mitochondrial parameters (right) in HepG2 cells after 24 h exposure. Data (at least four technical replicates) are expressed as mean ± standard deviation. Mean values with different symbols (*: compared to control; #: compared to MIX) within each mitochondrial parameter indicate significant differences (p < 0.05) among different treatments according to the one-way ANOVA test followed by the Tukey HSD multiple-comparison test as a post hoc analysis.

### 3.3. Impact of AFB1, FB1 and their combination on HepG2 transcriptomic responses

#### 3.3.1. Expression profiles of mRNAs in experimental groups

A total number of 29744 genes were detected after the exposure of HepG2 cells to either AFB1 (8 µg/mL) or FB1 (160 µg/mL) or their binary combination (MIX: 8 µg/mL for AFB1 and 160 µg/mL for FB1). A heatmap based on color key for the gene clustering is depicted in **Figure 5a**. Replicas from the same condition always cluster together, thereby generating four clusters according to HepG2 cell treatments. Regardless the treatment condition, the untreated control group had a distinct separation. The FB1 and MIX groups clustered together, thereby indicating a strong contribution of FB1 to the overall MIX effect compared to AFB1 alone, which yielded a clear distinct expression pattern compared to the other conditions (**Figure 5a**). Volcano plot based on the log Fold Change (FC) and the False Discovery Rate (FDR) of each tested gene is shown in **Figure 5b**. The cutoff value for the FDR was adjusted at 0.05, while log2FC < −1 for the downregulated genes and log2FC > 1 for upregulated genes were set to check the top significant genes. In comparison to the MIX group, AFB1 treated samples showed a large number of genes that downregulated (in the blue color) and upregulated (in red color). On the other hand, FB1 treated group had much less fold changes in the expressed gene in comparison to the MIX group (i.e. less number of the significant genes). The Venn diagrams in **Figure 5c** show the differential genes that are either up- or downregulated between the treatments. Compared to the untreated control (CON) scores, AFB1 resulted in 2336 different differentially expressed genes (DEGs), of which 558 DEGs are identical to those differentially expressed upon MIX treatment, which contained 312 upregulated DEGs and 246 downregulated DEGs. Similarly, compared to CON, FB1 treatment resulted in 2321 DEGs of which 1318 DEGs were identical to those differentially expressed upon MIX treatment, including 609 upregulated DEGs and 709 downregulated DEGs. Compared to the MIX condition, only 72 upregulated and 10 downregulated DEGs were shared between AFB1 and FB1, thereby confirming the different mode-of-action of both mycotoxins. In these Venn diagrams, it is also clearly visible that the single AFB1 condition is more different than the FB1, and thus contributes less to the MIX effects.

**Figure 5.**
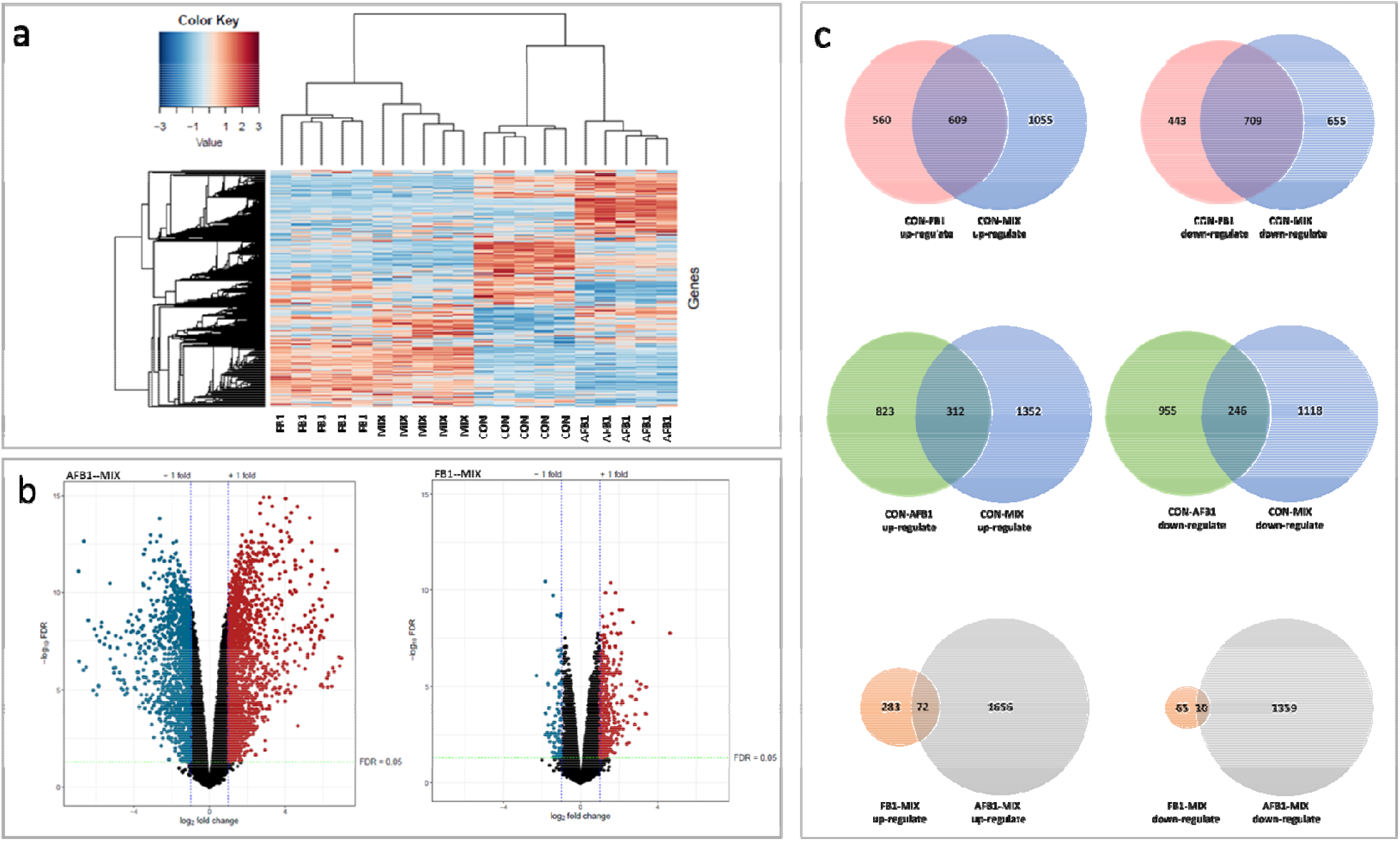
Differentially expressed genes in experimental groups. (a) Heatmap for outcoming of the weighted gene co-expression network analysis (WGCNA) on all conditions; (b) Volcano plot based on the fold change and the false discovery rate (FDR) of each tested gene. The cutoff for FDR was set at 0.05. Blue dots represent downregulated genes (log2FC < −1), red dots represent upregulated genes (log2FC > 1), black dots represent the genes that did not pass the thresholds for FDR and Log Fold Change; (c) Venn diagrams of the overlapping and different differentially expressed genes (DEGs).

#### 3.3.2. Kyoto encyclopedia of genes and genomes (KEGG) analysis-p53 pathway

Pathway analysis based on Gene Ontology (GO) (50 pathways) and Kyoto Encyclopedia of Genes and Genomes (KEGG) (338 pathways) databases led to the discovery of significantly enriched pathways upon AFB1 and FB1 treatment versus a combination of the two mycotoxins (MIX) treatment. Upon KEGG analysis, 6 significantly different KEGG pathways were identified when comparing FB1 and MIX (Herpes simplex virus 1 infection, Ribosome, Fanconi anemia pathway, Amyotrophic lateral sclerosis, Cell cycle, and p53 signaling pathway), whereas no differential pathways were identified between AFB1 and MIX. Overall these pathways, p53 signaling pathway is actively involved in bioenergetics and cell death. In **Figure 6**, it was shown the p53 signaling pathway genes that are differentially expressed with the comparison of FB1 and MIX. In this signaling pathway, MIX significantly downregulated Fas, DR5, Nora, PUMA, PIGs, Pag608 and upregulated p53, Bcl-xL, and Scotin. Subsequently, genes with mitochondrial and hence bioenergetic impact were significantly downregulated by MIX, including Cx I, Cx II, Cx III, Cx IV, PINK, and Bad. Cx I, Cx II, Cx III, and Cx IV are major mitochondria respiratory complexes in the electron transport chain (ETC), is linked to the CytC inducing apoptosis. As a result, further downregulated CytC, Apaf-a, CASP9, and CASP3, which may explain the cell death, possibly mediated by apoptosis in HepG2 cells. Because these differences were seen at the complete pathway level, there is a strong evidence that they pinpoint the major mode of action explaining the previous results.

**Figure 6.**
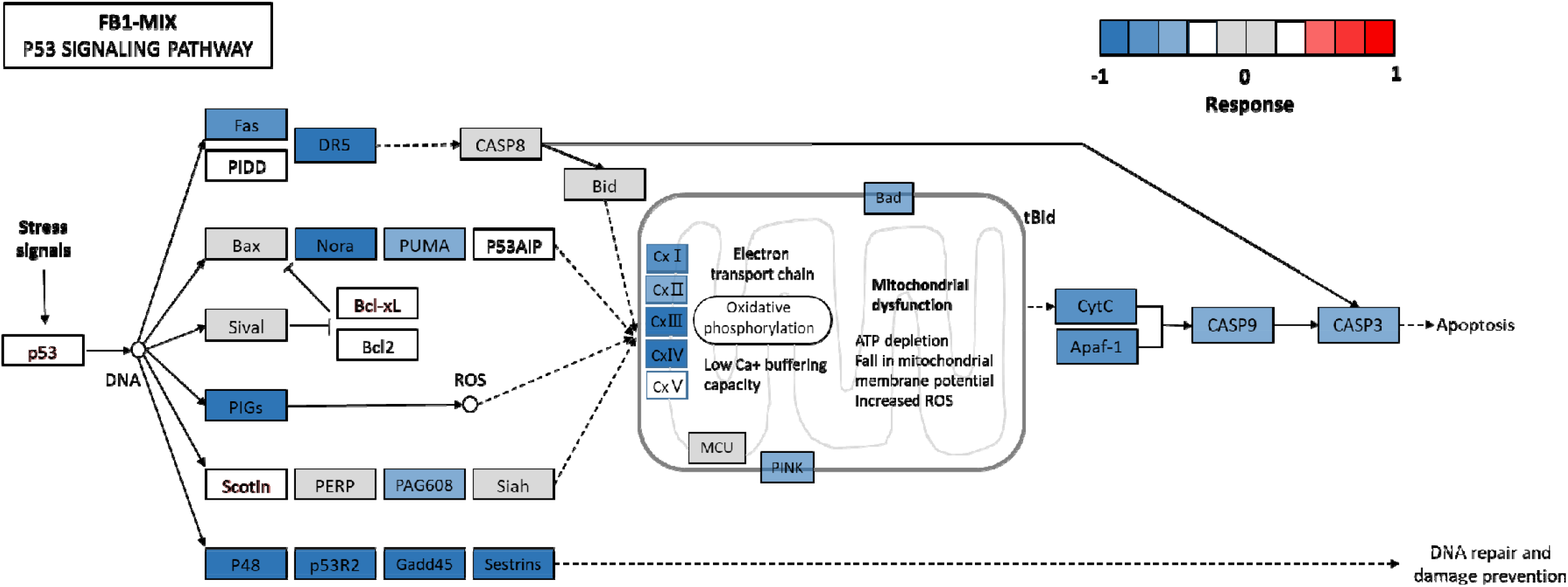
A pathway diagram of the p53 pathway of FB1-MIX as annotated by the Kyoto Encyclopedia of Genes and Genomes (KEGG). The color in which the genes are marked correlates to the response value of the comparison between fumonisin FB1 (FB1) and the AFB1-FB1 mixture (MIX).

#### 3.3.3. Gene ontology (GO) analysis

Gene Set Enrichment Analysis (GSEA) using the Hallmark gene sets showed that AFB1, FB1, and their binary combination (MIX) treatments affect key cellular processes. Key regulatory pathways (i.e. hypoxia, unfolded protein response, and p53 pathway), metabolic mechanisms (i.e. cholesterol homeostasis, OXPHOS, and glycolysis), and immune responses (i.e. TNF-α signaling via NF-κB, mTORC1 signaling, IFN-γ, IFN-α, and inflammatory response) were enriched in AFB1, FB1, and MIX versus CON, and in AFB1 and FB1 versus MIX. A heat map of z-scores of the Normalized Enrichment Score (NES) for each hallmark represents the differences in the enrichment of each hallmark between the pre-treatments (**Figure 7**). Upon GO analysis, seven differential pathways were identified when comparing FB1 and MIX (UV response DN, E2F targets, G2M checkpoint, p53 pathway, MYC targets V1, DNA repair, and mTORC1 signaling), and 4 differential pathways when comparing AFB1 and MIX (p53 pathway, MYC targets V2, DNA repair, and UV response DN). It was observed that AFB1 and FB1 mainly disrupted cell proliferation and as a result, MIX significantly altered proliferation genes compared to CON/AFB1/FB1. In addition, AFB1 also showed significant induction of DNA damage.

**Figure 7.**
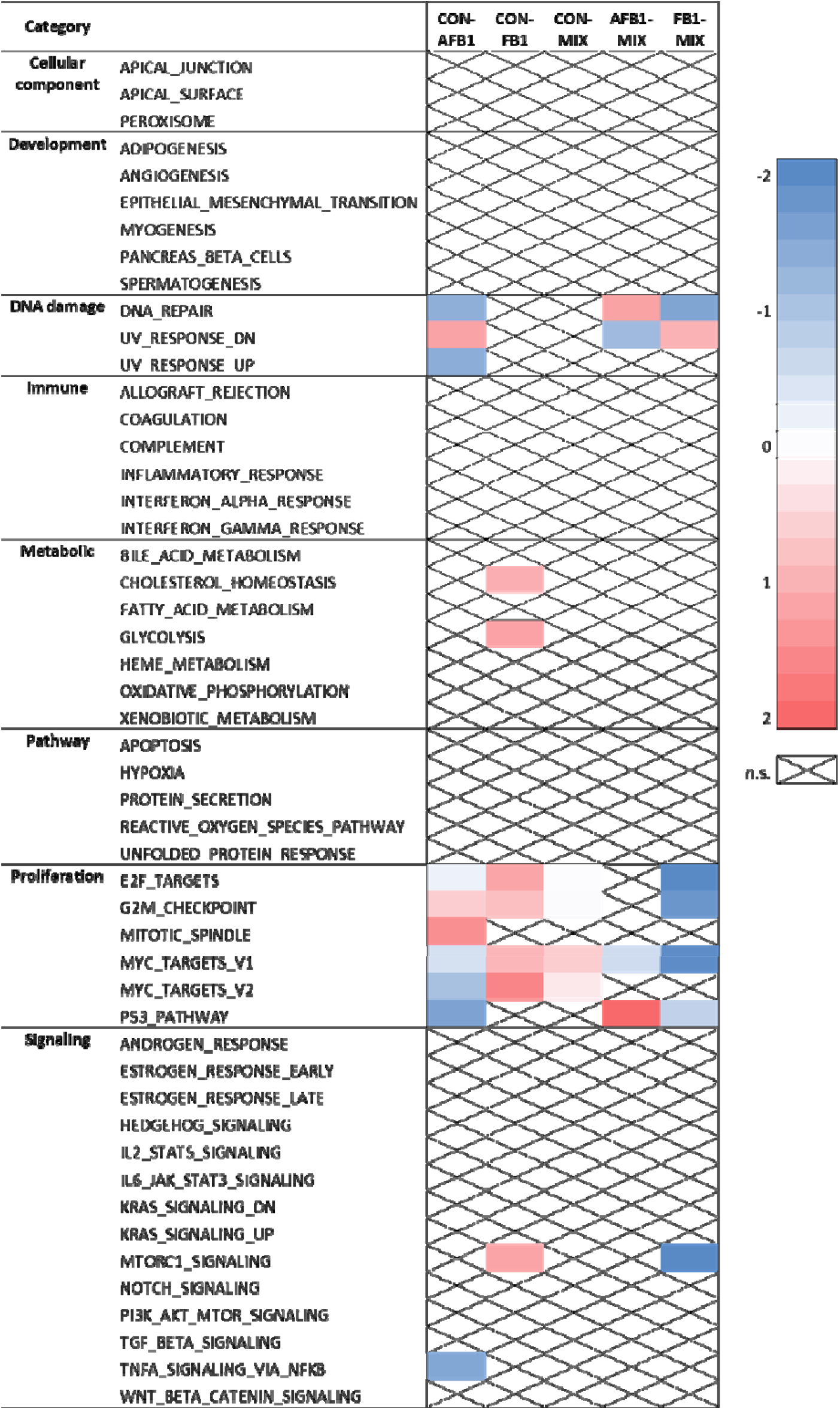
Heatmap of Z-scores of the MSigDB hallmark gene sets using Gene Set Enrichment Analysis (GSEA), showing the significantly enriched gene sets (FDR<0.05) across the treatments according to a pairwise comparison. Non-significant (n.s.) differences are marked by X.

## 4. Discussion

The current study aimed at identifying the short term effect of (combined) exposure of AFB1 and FB1 on the cellular energy profiles, including glycolysis and mitochondrial respiration pathways, in HepG2 cells. This was accomplished by applying relevant doses and combinations of both mycotoxins in which the low level of exposure or treatment is matching with average of urinary biomarkers of AFB1 (0.5 µg/mL) and FB1(10 µg/mL) in humans (Meneely et al. 2018). Two more levels of exposure were applied by increasing the applied doses four folds to have a middle exposure scenario and eight folds as high exposure scenario to investigate the potential toxicity. It was found that with advanced respirometry techniques, differential effects between a single mycotoxin treatment compared to the binary mixture became obvious in multiple mitochondrial parameters. Finally, transcriptomics clearly showed distinct effects amongst all different treatments, and differential pathways and genes revealed a particular focus on mitochondrial and proliferation-related mechanisms.

### 4.1. Conventional assays confirm earlier findings on cytotoxicity, oxidative stress, and mitochondrial membrane potential

AFB1, FB1, and their binary mixture (MIX) individually caused cytotoxic effects, ROS generation, and MMP disruption in HepG2 cells, and MIX shows a higher toxic effect for these cytotoxicity endpoints. It is presumed that MIX may aggravate mitochondrial dysfunction, resulting in an increase in ROS generation and induction in MMP disruption. Previous research elucidated the biochemical principles that drive mitochondrial respiration, including the tricarboxylic acid (TCA) cycle and fatty acid β-oxidation in the mitochondrial matrix, that generate electron donors to fuel respiration, ETC complexes, and ATP synthase in the inner mitochondrial membrane (IMM) carrying out oxidative phosphorylation (Vyas et al. 2016). MMP is generated by proton pumps (complex I, III, and IV) through ETC (Zorova et al. 2018). ROS is generated by incomplete electron transfer through ETC complexes I and II resulting in O^2−^ production in the mitochondrial matrix, while an electron leak at complex III generates O^2−^ in both matrix and intermembrane space (Feissner et al. 2009). However, when comparing the effects of AFB1/FB1 and MIX in each applied condition, MIX did not significantly modify cytotoxicity endpoints compared to the individual toxins (ROS and MMP), which may be caused by the different cytotoxicity-related mode-of-action of AFB1 and FB1 in HepG2 cells. Indeed, the toxicology of AFB1 is intimately linked with its biotransformation to the highly reactive AFB1-exo28,9-epoxide (AFBO), which produces direct genotoxicity through the formation of adducts with the DNA, and to a lesser extent the induction of oxidative stress, which is responsible (probably among other causes) for the indirect genotoxicity of AFB1 (Zhu et al. 2021). This mode of action was confirmed with the transcriptomics outcomes, in which AFB1-containing treatments involve DNA damage pathways. On the other hand, FB1 could cause liver toxicity and the most recognized mechanism of action is the disruption of sphingolipid metabolism by inhibiting the ceramide synthase enzyme (Abdul and Chuturgoon 2021).

### 4.2. Respirometry reveals interactions between mycotoxins at the level of the mitochondria

To investigate whether the biological processes such as total ATP production, glycolytic and mitochondrial function are affected by the AFB1/FB1 and MIX, their cellular rates using the Seahorse XF96 instrument were analyzed. The main metabolic routes contributing to energy homeostasis are glycolysis and OXPHOS, which couple the breakdown of nutrients such as glucose, amino acids, and fatty acids to ATP production (Fox et al. 2005). These two pathways also play a pivotal role in redox homeostasis since they contribute to the reducing power required for anabolic processes (Fox et al. 2005). The Seahorse XF Real-Time ATP Rate Assay allows the calculation of the mitochondrial and glycolytic ATP production rates, which provides a new dynamic and quantitative insight into cellular bioenergetics by providing a real-time measurement of oxygen production as a proxy for respiration, and lactate secretion as a proxy for glycolysis. The Seahorse XF Glycolysis Stress Test and Mitochondrial Stress Test protocols dissect the glycolytic and respiratory fluxes components into basal, maximal, and reserve (spare) glycolytic or respiratory capacity through the consecutive addition of the stressors such as oligomycin, FCCP, and rotenone in HepG2 cells, whose optimization is reported in **Figure 2, 3,** and **4**.

AFB1, FB1, and MIX individually disrupted ATP production by glycolysis and mitochondrial respiration or OXPHOS, which is in line with the cytotoxicity data. Interestingly, the MIX condition showed a higher interference with total ATP production metabolism in HepG2 cells compared to the single mycotoxin treatments. In addition, MIX shifted the fraction of ATP production between OXPHOS and glycolysis from 43 %/57 % to 67 %/33 % under high condition, thereby indicating a particular decrease in glycolysis. Generally, cellular metabolism consumes energy of which 70 % is supplied by OXPHOS, although cell type type-dependent differences are reported (Zheng 2012). As HepG2 cells have a cancer-derived origin, energy metabolism and glucose and glutamine uptake are different compared to normal tissues and display a high rate of glycolysis (Zheng 2012). Due to their different origin and differentiation, glycolysis contributes to most of ATP but does not generally exceed 50–60 % in cancer cells (Zu and Guppy 2004). Therefore, according to our data, it is inferred that combined AFB1 and FB1 could suppress energy metabolism and change metabolic phenotype to adapt to microenvironmental changes, which may result in a selective advantage for HepG2 cells to survive under an unfavorable environment (Marusyk and Polyak 2010). Tumor cell proliferation requires sufficient metabolic flux through the pentose phosphate pathway (PPP) to meet the demand for biosynthetic precursors and to increase protection against oxidative stress, which in turn requires upregulation of substrate flow through glycolysis. This metabolic poise is often coupled with a shift in ATP production from mitochondrial OXPHOS to substrate-level phosphorylation (Skolik et al. 2021). The PPP branches from glycolysis at the first committed step of glucose metabolism, which is catalyzed by hexokinase and consumes glucose-6-phosphate (G6P) as a primary substrate. It is required for the synthesis of ribonucleotides and is a major source of NADPH, which is required for and consumed during fatty acid synthesis and the scavenging of ROS (Skolik et al. 2021). Therefore, the PPP plays a pivotal role in helping glycolytic cancer cells to meet their anabolic demands and combat oxidative stress (Patra and Hay 2014). In the PPP, Glucose-6-phosphate dehydrogenase (G6PD) is an enzyme that catalyzes the first reaction, providing reducing power to all cells in the form of NADPH (Patra and Hay 2014). Therefore, the damage of G6PD may hinder or slow the supply of energy through glycolysis. In our study, it is shown that the fraction of total ATP production started to be shifted in HepG2 cell exposure to middle concentration of AFB1, and then there is a significant shift in HepG2 cell exposure with all high concentrations of MIX. It is hypothesized that AFB1 may reduce the G6PD activity to inhibit glycolysis to produce ATP, which leads to this shift of ATP in HepG2 cells. Raafat et al. have reported that AFB1 exposure is associated with the evident decline in the activity of G6PD enzyme (Raafat et al. 2021). In addition, Liu et al. also mentioned that there may be a novel association of G6PD activity with AFB1-related xenobiotic metabolism (Lin et al. 2013). These previous studies could support our conjecture that AFB1 may reduce the G6PD activity. Our previous studies showed that other microbial toxins, namely *Bacillus cereus* emetic toxin cereulide induces toxicity, which relies on the mitochondrial dysfunction in HepG2 cells. Oxygen consumption rate analyses and the bioenergetics assessment with Seahorse XF analyzer showed measurable mitochondrial impairment at doses of cereulide even lower than here used AFB1 and FB1. Observed mitochondrial dysfunction was greatly reflected in reduction of maximal cell respiration.

When considering the specific glycolytic and mitochondrial parameters individually, it was observed that AFB1, FB1, and MIX can significantly inhibit HepG2 cells to produce ATP in both pathways. Especially upon exposure to high mycotoxin concentrations, MIX showed a significant decrease in mitochondrial respiration with all mitochondrial parameters (basal respiration, maximal respiration, ATP production, proton leak, spare respiration capacity, and non-mitochondrial respiration) in HepG2 cells compared to only-AFB1/-FB1, while no significance in glycolysis parameters were observed between MIX and single. A decrease in the ATP-linked OCR may indicate a low ATP demand, a lack of substrate availability, and/or severe damage to OXPHOS. The remaining rate of mitochondrial respiration is defined as the proton leak, and consist of protons transported through the mitochondrial membrane during electron transport, that result in oxygen consumption but not ATP production. The spare respiratory capacity (SRC) characterizes the mitochondrial capacity to meet extra energy requirements, beyond the basal level, in response to acute cellular stress or heavy workload and thereby avoiding an ATP crisis, and can be viewed as a determination of mitochondrial fitness, a reflection of “healthy” mitochondria (Marchetti et al. 2020). When cells are subjected to stress, the energy demand increases, with more ATP required to maintain cellular functions (Yamamoto et al. 2016). Our results demonstrated that the combination of AFB1 and FB1 probably leaded to a significant interaction by causing more disruption of the mitochondrial metabolism resulting in apoptosis (involving complex I-V). As a mode of action, AFB1 exposure can also cause hepatotoxicity at the DNA level, which is accompanied by several metabolic changes including cell membrane metabolism, glycolysis, and TCA cycle functioning, and mainly cause oxidative-stress-mediated impairments of mitochondria functions (Zhang et al., 2011; Zhou et al., 2021). In line with previous research, AFB1 impairs mitochondrial respiration, causes MMP loss, reduces ATP content, and inhibits the function of mitochondrial complexes I-IV (Chen et al. 2022; Xu et al. 2021). Similarly, FB1 is involved in mitochondrial dysfunction by inhibiting ETC in mitochondrial respiration (Chen et al. 2022; Sheik Abdul and Marnewick 2020). Therefore, AFB1 and FB1 could disrupt mitochondrial respiration by ETC, which could be the reason that the mixture of AFB1 and FB1 worsened the mitochondrial dysfunction and showed a significant interaction in the disruption of the mitochondrial metabolism. As mitochondrial respiration is affected in many pathologic conditions such as hypoxia and intoxications, the impaired electron transport chain could emit additional p53-inducing signals and thereby contribute to cell damage (Khutornenko et al. 2010). Our transcriptomic results and literature report that AFB1 and FB1 increase p53 expression, which could be another reason for a significant interaction between AFB1 and FB1 on mitochondria damage (Cao et al. 2022; Molina-Pintor et al. 2022).

### 4.3. Transcriptomics reveals interactions between both mycotoxins at the level of mitochondrial functioning and apoptosis

Both metabolic flux measurements from seahorse assays and RNA sequence analysis of AFB1, FB1, and MIX in HepG2 cells indicated mode-of-actions related to cell death, apoptosis, or mitochondrial dysfunction. Interestingly, it was observed that MIX has more similar DEGs with FB1 than with AFB1, thereby suggesting that FB1 dictates the combined response more than AFB1. Nevertheless, compared to AFB1 and FB1 together, MIX could upregulate 72 DEGs (CPLX2, DDX46, ABCC11, SARDH, and CYP24A1 genes: related to mitochondrial metabolism) and downregulate 10 DEGs (RFX2 gene: DNA-binding protein; CD274 gene: programmed cell death), which suggests that AFB1 and FB1 may co-regulate the expression of some genes resulting in a significant interaction. Our results showed that the p53 pathway is one of the co-regulated signaling pathways by AFB1 and FB1, and is related to mitochondrial dysfunction resulting in apoptosis. The p53 pathway is a major orchestrator of the cellular response to a broad array of stress types by regulating mitochondrial apoptosis, DNA repair, and genetic stability, and participates directly in the intrinsic apoptosis pathway by interacting with the multidomain members of the Bcl-2 family to induce mitochondrial dysfunction (Vaseva and Moll 2009). This system is essential in humans for genome integrity, DNA repair, and apoptosis (Bernstein et al. 2002).

In apoptosis, the stabilization and activation of p53 lead to programmed cell death (Yu et al., 2009), where p53 directly upregulates the expression of cell surface death receptors proteins such as Fas/APO1 and KILLER/DR5 (Burns and El-Deiry 1999). Cytoplasmic pro-apoptotic proteins like PIDD and Bid are also thought to be the putative target of p53 (Sax and El-Deiry 2003). In addition, mitochondrial pro-apoptotic proteins such as Bax, Bak, PUMA, and NOXA are also regulated by p53 (Oda et al. 2000; Wei et al. 2001; Yu et al. 2003). In this study, Fas, DR5, PUMA, Noxa, and PIGs genes were significantly downregulated by MIX compared to single FB1 exposure, which could be associated with mitochondrial dysfunction. It also has shown that MIX significantly upregulated p53 gene and downregulated Cx I, Cx II, Cx III, and Cx IV genes comparing single FB1 exposure in the p53 signaling pathway, which could be the reason for a significant interaction between AFB1 and FB1 on mitochondrial damage. As mitochondrial respiration is affected in many pathologic conditions such as intoxications, the impaired ETC could emit additional p53-inducing signals and thereby contribute to tissue damage (Khutornenko et al. 2010). It is also reported that a strong p53 response is induced specifically after inhibition of the mitochondrial cytochrome bc1 (the electron transport chain complex III). Accumulation of p53 in the mitochondria is observed in animal and cell culture models and is associated with mitochondrial depolarization and mitochondrial complex IV inactivity (Marchenko and Moll 2014). Saleem et al. also mentioned that lower complex IV activity and several impaired indexes of mitochondrial function are related to p53 (Saleem et al. 2015). AFB1 and FB1 individually affected the p53 and complexes, which could result in the MIX significantly upregulated p53 gene and downregulated Cx I, Cx II, Cx III, and Cx IV genes resulting in the significant inhibition interaction on mitochondrial respiration. As reported in the literature, HepG2 cells exposed to either AFB1 or FB1 showed a higher abundance of p53 (Budin et al. 2021; Li et al. 2021). Moreover, Du et al have shown that when AFB1 and FB1 were combined, a higher optical density of p53 was observed by immunohistochemical analysis, and they hypothesized that there could be an interaction between AFB1 and FB1 in inducing HepG2 cell apoptosis (Du et al. 2017). This finding is consistent with our results as in the current work MIX also significantly upregulated p53.

It has been verified that AFB1 inhibits mitochondrial complexes I-IV activities and FB1 inhibits mitochondrial complex I by decreasing complex sphingolipids (Huang et al. 2020). This is also in a line with our results, where MIX resulted in a significant downregulation of Cx I, Cx II, Cx III, and Cx IV genes. These genes are major mitochondria respiratory complexes in ETC, and linked to cytochrome c (CytC) inducing apoptosis. CytC, a role in cell apoptosis in p53 pathway, is released into the cytosol, and then the protein binds to Apaf-1, activates CASP9, and triggers an enzymatic cascade leading to cell death (Schuler et al. 2000). In our study, the expression of CytC, Apaf-1, CASP9, and CASP3 genes was decreased by MIX compared to FB1. The release of CytC from mitochondria is a central event in the death receptor-independent, “intrinsic,” apoptotic pathway (Desagher and Martinou 2000). CytC is essential for the assembly and respiratory function of the enzyme complex, and the lack of CytC decreases the stability of complex IV, reduces electron transport complex III activity, and modifies redox metabolism (Welchen et al. 2012). CytC together with ATP and Apaf-1 facilitates activation by CASP9 of the effector caspases CASP3 (Slee et al. 1999), which then cleaves their substrates, finally leading to the apoptotic cell death. This complex of CytC, Apaf-1 and CASP9 is commonly referred to as the apoptosome (Bratton et al. 2001). The reduced form of CytC also binds less to anions and binds less tightly to negatively charged membranes. This could be the reason for a significant interaction between AFB1 and FB1 on mitochondria dysfunction and HepG2 cell apoptosis by disrupting the mitochondrial complexes and CytC in p53 pathway.

DNA repair is another system in the p53 pathway. The stabilization and activation of p53 lead to cell cycle arrest by increasing GADD45 (Jin et al. 2002), initiating DNA repair through p53R2 and p48 (Tanaka et al. 2000). Cells that are defective in DNA repair tend to accumulate excess DNA damage. In addition, cells defective in apoptosis tend to survive even with DNA damage, and the subsequent DNA replication during cell division may cause persistent mutations leading to carcinogenesis (Bernstein et al. 2002). Normally, DNA damage is repaired by base excision repair (BER) by mitochondrial enzymes. Mitochondrial DNA comprises 0.1-1.0 % of the total DNA in most mammalian cells (Singh et al., 1992). Mitochondrial DNA has been proposed to be involved in carcinogenesis because of its high susceptibility to mutations and limited repair mechanisms in comparison to genomic DNA (Penta et al. 2001). Mitochondrial DNA damage, if not repaired, leads to disruption of the ETC and mitochondrial dysfunction (Mandavilli et al. 2002). In general, the energy-demanding process of DNA repair is the proper utilization of the available ATP in the cell which is provided by the mitochondria (Bernstein et al. 2002). Therefore, mitochondrial DNA repair plays a central role in maintaining (energy) homeostasis in the cell. In our study, P48, p53R2, Gadd45, and Sestrins genes were significantly downregulated by MIX compared to FB1 treatment in HepG2 cells. This suggests that the combination of AFB1 and FB1 could have a significant inhibition interaction on the DNA repair system and thus cell homeostasis.

Our study also shows that AFB1, FB1, and MIX disrupted HepG2 cell proliferation, by means of E2F targets, G2M checkpoint, mitotic spindle, MYC targets, and p53 pathway. Deregulated cell proliferation could propel the tumor cell and its progeny into uncontrolled expansion and invasion beneath the complexity and idiopathy of every cancer. Neoplastic progression could be further supported by the deregulated cell proliferation that is, together with the obligate compensatory suppression of apoptosis, needed to support it (Evan and Vousden 2001). Previous studies have illustrated that individual AFB1 and FB1 had an ability to induce proliferation to increase apoptosis (Singh and Kang 2017; Zhou et al. 2019). Therefore, we speculate that the combination of AFB1 and FB1 may deregulate proliferation, as a result, triggering apoptosis.

## 5. Conclusion

The cytotoxicity of AFB1 and FB1 to HepG2 cells has been examined from cytotoxicity endpoints (cell viability, ROS generation, and MMP disruption), total ATP production, glycolytic, mitochondrial function, and gene expression in the cell apoptosis process. The combined exposure of both mycotoxins induced more inhibitory effect on the cellular viability, an increase in the ROS production, and a disruption of MMP. Respirometry and transcriptomics demonstrated a significant interaction between AFB1 and FB1 in pathways related to mitochondrial dysfunction and apoptosis, which is most probably triggered by the p53 pathway and mitochondrial complex Cx I-IV genes. In addition, AFB1 and FB1 affected DNA repair and induce cell proliferation in HepG2 cells in a possible synergistic nature, because of their different targets in cell apoptosis.

## Abbreviations

AFB1: Aflatoxin B1
FB1: fumonisin B1
MIX: their binary mixture
CON: control
HepG2: human hepatocellular carcinoma
IARC: International Agency for Research on Cancer
ATP: adenosine triphosphate
ROS: reactive oxygen species
DMEM: Dulbecco’s modified Eagle’s medium
NEAA: non-essential amino acids
FBS: Fetal Bovine Serum
PBS: Phosphate buffer saline
MTT: tetrazolium salt
MMP: mitochondrial membrane potential
OXPHOS: oxidative phosphorylation
ECAR: extracellular acidification rate
OCR: oxygen consumption rate
DEGs: differentially expressed genes
FC: fold change
FDR: false discovery rate
GO: Gene Ontology
KEGG: Kyoto Encyclopedia of Genes and Genomes
ETC: electron transport chain
IMM: the inner mitochondrial membrane
AFBO: AFB1-exo28,9-epoxide
PPP: pentose phosphate pathway
G6P: glucose-6-phosphate
G6PD: glucose-6-phosphate dehydrogenase.

## CRediT authorship contribution statement

**Xiangrong Chen:** Conceptualization, Methodology, Validation, Formal analysis, Investigation, Resources, Data curation, Writing – original draft. **Mohamed F. Abdallah**: Formal analysis, Data curation, Writing - Review & Editing. **Charlotte Grootaert**: Conceptualization, Validation, Data curation, Writing - Review & Editing. **Filip Van Nieuwerburgh**: Data curation, Methodology, Writing - Review & Editing. **Andreja Rajkovic**: Conceptualization, Methodology, Project administration, Funding acquisition, Writing – review & editing, Supervision.

## Acknowledgments

The authors wish to thank China Scholarship Council (CSC) for providing X.C. with a full personal (not including bench fee) Ph.D. scholarship (File No. 201806170042) to study at Ghent University, Belgium. M.F.A is supported by the Ghent University Special Research Fund (BOF) postdoc mandate with grant number BOF20/PDO/032. Authors and AR in particular express gratitude to European Commission for the received funding for this research performed as part of ImpTox project (grant agreement No <u>965173)</u>, Research Foundation Flanders for the Research grant provided to AR (1506419N), and Ghent University Special Research Fund given to AR for purchase of Seahorse XF analyzer (BOF20/BAS/120). Authors thank Ghent University, and their respective Faculties and Departments for team work, support and general infrastructure.

## Data availability

All data generated or analyzed in this study that are relevant to the results presented in this article are included in the article.

